# Long-term rapamycin treatment suppresses IL-17-producing gamma delta T cells and blunts neuroinflammation in aging

**DOI:** 10.64898/2026.02.04.703808

**Authors:** Clement Torrent, Caterina Gagliardi, Nina Fülle, Ignazio Antignano, Maria Eugenia Bernis, Miriam Stork, Daniele Bano, Melania Capasso, Lily Keane

## Abstract

Aging is the gradual accumulation of structural and functional changes in an organism over time, including immune remodeling and a progressive increase in basal inflammation, or inflammaging. The mTOR pathway is a central driver of aging-related diseases, such as cancer, chronic inflammation and neurodegeneration; pharmacological inhibition with rapamycin is associated with reduced aged-related morbidity and increased lifespan across species. Nonetheless, concerns remain about the use of rapamycin, a well-established immunosuppressant in transplant medicine, as an anti-aging intervention. Here, we evaluated the impact of prolonged low-dose dietary rapamycin on the aging immune system. Treatment did not significantly alter innate or adaptive immune cell populations, including brain resident microglia; however, it attenuated the age-associated accumulation of IL-17–producing γδ T cells, particularly in the peritoneal cavity. After a peripheral inflammatory LPS challenge, circulating IL-17 levels were significantly reduced and correlated with an attenuation of microglia inflammatory phenotype. These findings suggest that prolonged low-dose rapamycin exposure exerts minor systemic immune changes, while selectively limiting age-related γδ T cell expansion and neuroinflammation associated with systemic inflammation.

## Introduction

Aging is a process characterized by the progressive manifestation of hallmarks such as stem cell exhaustion, cellular senescence, chronic inflammation and metabolic dysfunction, leading to the loss of physiological homeostasis and increasing risk of death (1, 2). The mechanistic target of rapamycin (mTOR) complex 1 (mTORC1) pathway fine-tunes a majority of these aging hallmarks (3). Therefore, significant attention has been dedicated to understanding the role of the mTORC1 pathway in aging.

The serine-threonine kinase mTOR is activated by a broad range of stimuli, such as growth factors, cytokines and amino acids (4-6) and, when associated with other proteins to form mTOR complex 1, it integrates catabolic and anabolic signals to balance cell metabolism, survival, growth, proliferation, migration, differentiation and immune response (6). Given its critical role in regulating cell responses, mTORC1 activity is tightly linked with several age-related pathologies, such as neurodegenerative diseases, cancers, diabetes, and chronic inflammation (7).

In the past two decades, numerous studies have investigated the potential of mTORC1 inhibitors as treatment for several cancers and as immunosuppressants (8-10). One of these inhibitors is rapamycin, the first TOR inhibitor to be identified (11). Rapamycin or Sirolimus has been widely used in transplantation for its immunosuppressive properties (12). Paradoxically, short-term rapamycin treatment has also been shown to enhance vaccine responses, an effect attributed in part to heightened myeloid cell activity (13-16). Beyond its immunomodulatory roles, renewed interest in rapamycin and mTORC1 inhibition was sparked by the discovery that rapamycin extends lifespan (17, 18), an effect thought to be mediated largely through suppression of cancer development (8). Nonetheless, while accumulating evidence suggests that rapamycin ameliorates certain aspects of age-associated inflammation, its overall impact on the aging immune system remains incompletely understood. Notably, literature remains elusive on how rapamycin modulates acute systemic inflammatory responses in aging, for example responses to systemic LPS. For these reasons, doubts remain on whether long-term *in vivo* rapamycin treatment, and therefore mTORC1 inhibition, would be a safe anti-aging approach or cause excessive immunosuppression. Indeed, in rats, rapamycin inhibited wound healing (19) and in Tuberous Sclerosis Complex patients, which have an aberrant hyperactivation of mTOR signaling, rapamycin led to an increase in dermatologic adverse events such as oral ulceration and acneiform eruptions (20).

In this study, we aimed to understand how long-term, low-dose rapamycin could affect the ageing immune system under both basal and inflammatory conditions. To this end, mice were given a low dose of dietary rapamycin for a period of 5 months and immune cell proportions and inflammatory markers were then measured at steady state and upon and LPS challenge. We found that long-term treatment did not substantially alter immune cell populations, nonetheless, it counteracted the age-related increase of IL-17 producing γδ T cells and moderately blunted peripheral and brain-related inflammation. Together, our data reveal that long-term rapamycin treatment could be a potential intervention to modulate age-associated IL-17-responses.

## Results

### Rapamycin does not alter myeloid cells skewing in aging

In order to assess the broad effect of long-term rapamycin exposure on aging of the immune system, we designed a study to compare mice fed a rapamycin-containing diet (samples named “Old Rapa”) vs mice fed with a diet containing only the encapsulating compound EUDRAGIT® (“Old Eud”). The diet composition and rapamycin dose were selected based on the original study by Zhang *et al*., demonstrating lifespan extension following rapamycin treatment (21). Mice were placed on the diet at 17-19 months and assessed 5 months later. Three months old mice fed a standard diet were analyzed simultaneously for comparison (**Fig 1A**).

**Figure 1.**
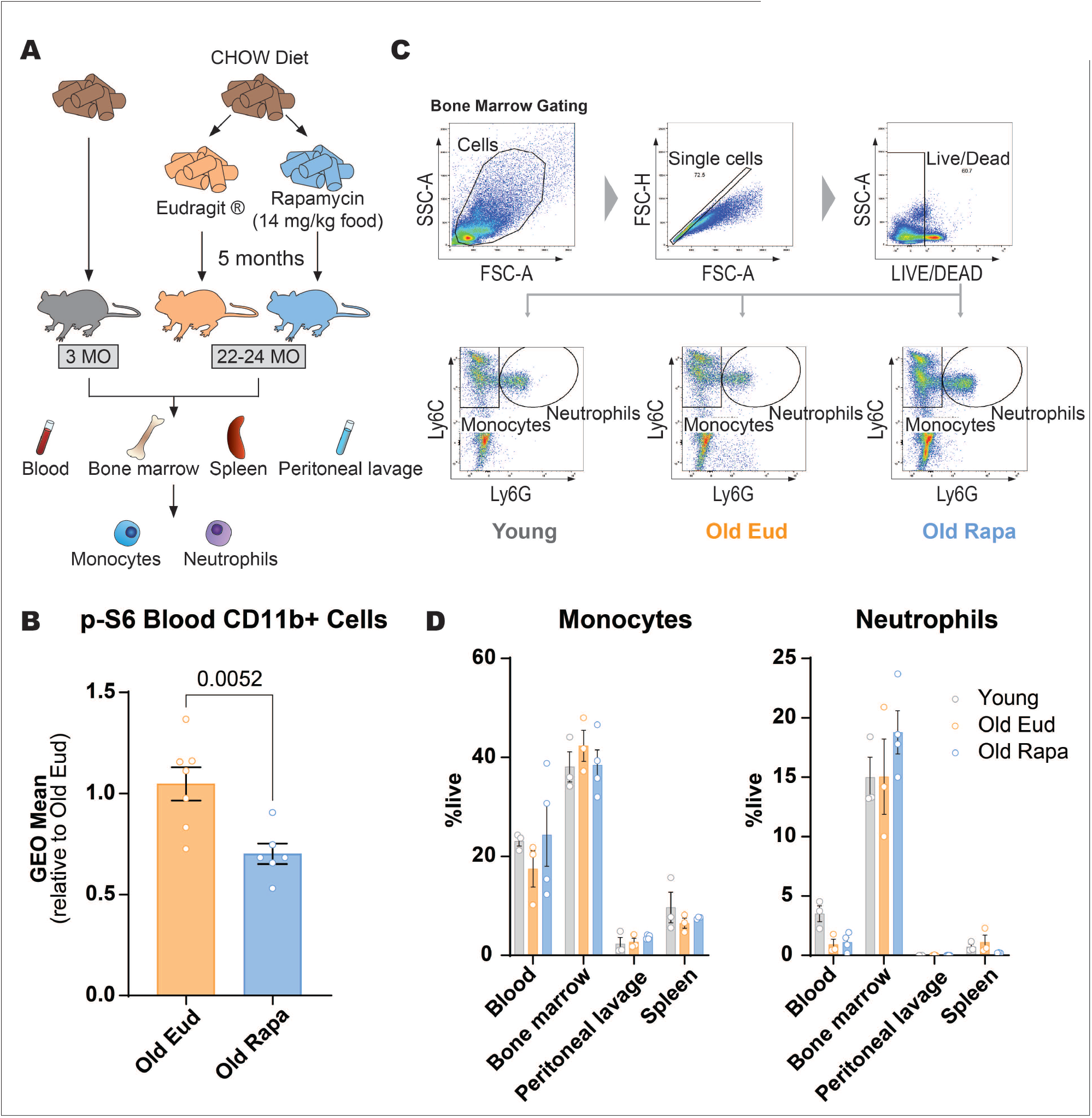
Long-term, low-dose rapamycin does not alter the proportion of myeloid cells. (**A**) Schematic model of young and old mice exposed to rapamycin diet with subsequent flow cytometry analysis of immune myeloid cells. (**B**) p-S6 levels in CD11b+ cells from blood of old Eudragit and rapamycin mice, pooled from two independent cohorts. The first cohort is represented in panel (**A**), in the second cohort, treatment was initiated at 8 months of age and mice were analyzed at 13 months. (**C**) Representative gating strategy for live cells expressing Ly6C and Ly6G to identify monocytes and neutrophils (n = 3–4 per group). (**D**) Monocytes and neutrophil populations from blood, bone marrow, peritoneal lavage and spleen from young, old Eudragit and old rapamycin mice, (n = 3–4 per group). Data are shown as mean ± s.e.m. Statistical analysis was conducted by Welch’s t-test (**B**) and One-way ANOVA (**D**).

Rapamycin-containing diet reduced mTORC1 signaling, as indicated by reduced phosphorylation of ribosomal protein S6 (p-S6) in CD11b+ myeloid cells in the blood. The decrease in p-S6 levels was comparable between two different cohorts, despite differences in age at the start of treatment (8 months vs. 17–19 months), underlying that a similar extent of mTOR inhibition can be achieved at this dose and treatment duration, irrespective of age **(Fig 1B)**.

In order to assess the effect of rapamycin on proportions of peripheral immune cells, we first assessed mature myeloid cells by flow cytometry. Previous publications have highlighted that aging is associated with alterations in immune homeostasis, notably, shifts to a higher myeloid cellular output also called “myeloid skewing” (22-23).

At the study endpoint, blood, bone marrow, spleen, and peritoneal lavage were collected and the percentage of Ly6C+/Ly6G– monocyte and Ly6C+/Ly6G+neutrophil populations was assessed (**Fig 1C**). Monocyte and neutrophil proportions were quantified as percentage of live cells.

In our cohort, we did not observe an effect of rapamycin on monocyte or neutrophils in any of the tissues investigated (**Fig 1D**) or an increase of the percentage of myeloid cells with age (**Fig 1D**).

These data collectively suggest that rapamycin does not exert any significant effect on monocytes and neutrophils numbers, as their percentage in blood, bone marrow, peritoneal lavage and spleen in aged mice was not affected.

### Rapamycin does not affect B cell populations in older mice

We next investigated if a rapamycin-containing diet could impact the lymphoid lineage, focusing first on B cells in the bone marrow, lymph nodes, peritoneal lavage, and blood (**Fig 2A**).

**Figure 2.**
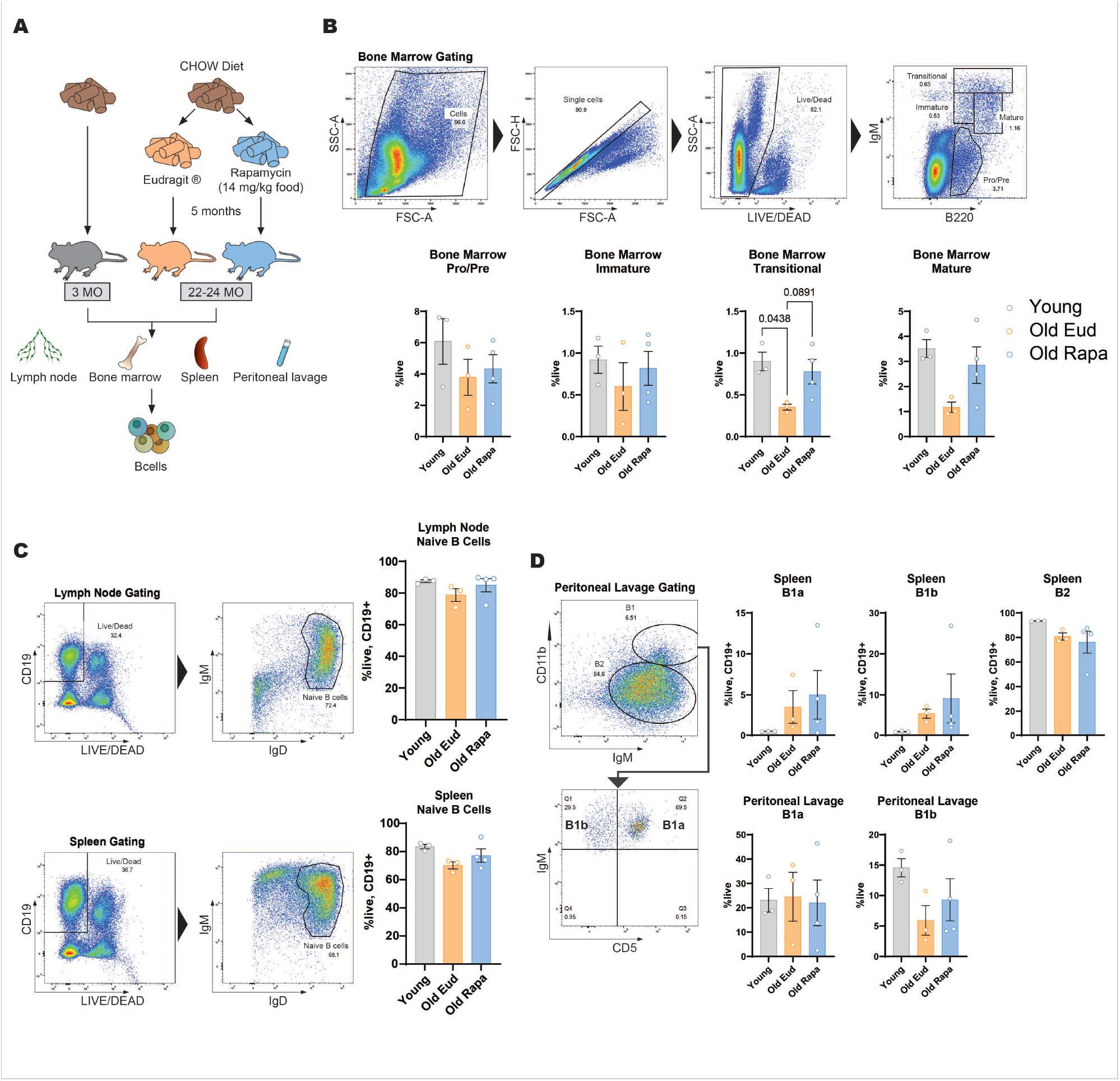
Rapamycin does not significantly alter B cell subsets, but shows a trend toward rescuing transitional and mature B cells in aged mice. (**A**) Schematic representation of the experimental setup. (**B**) Representative flow cytometry gating strategy for bone marrow B cell subsets and their quantification (n = 3–4 per group). (**C**) Representative gating strategy and quantification of naïve B cells from lymph nodes and spleen (n = 3–4 per group). (**D**) Representative gating strategy and quantification of B1 and B2 cells, including B1a and B1b subsets, from peritoneal lavage (n = 3–4 per group). Data are presented as mean ± s.e.m. Statistical analysis was conducted by One-way ANOVA.

B-cell differentiation in the bone marrow was assessed by measuring cells expressing B220 and the *μ*- chain of immunoglobulin M (IgM). No significant changes were observed in the proportions of pro-B cells, pre-B cells and immature B cells (**Fig 2B**). However, although not statistically significant, we noted a trend towards an increase in mature and transitional B cells in aged mice receiving rapamycin, resembling levels observed in young mice (**Fig 2B**).

Regarding naïve B cells in lymph nodes and spleen, no substantial differences were noticed across the three groups (**Fig 2C**).

Finally, we investigated B1 and B2 cell populations in the spleen and peritoneal lavage based on IgM, CD11b and CD5 markers. Consistent with previous findings, no significant differences were detected across B cell subpopulations, including subsets of B1 cells such as B1a and B1b, in the spleen and in the peritoneal lavage (**Fig 2D**).

Altogether, these results indicate that low-dose rapamycin administration has a minimal impact on B cells in aging.

### Rapamycin-treated mice show minor changes in naïve and memory T-cell populations

Aging is associated with a decline in naïve T cell numbers and an increase in memory T cells (24, 25). In previous studies, rapamycin has been shown to increase differentiation of memory CD8 T cells (26, 27), however, mTOR-deficiency impairs differentiation of CD4 T cells to Th1, Th2 and Th17 subsets (28). Therefore, we assessed whether rapamycin treatment could influence the age-associated skewing of naïve to memory T cell populations. To this end, we assessed naïve (CD62L+CD44–), central (CD62L+CD44+) and effector memory (CD62L–CD44+) CD4 and CD8 T cells by measuring the expression of lymph node homing receptor, CD62L, a marker for naïve and central memory T cells, together with CD44, a marker of T cell memory and activation. Helper (CD4+) and cytotoxic (CD8+) T cells were assessed from lymph node, spleen and peritoneal lavage (**Fig 3A**).

**Figure 3.**
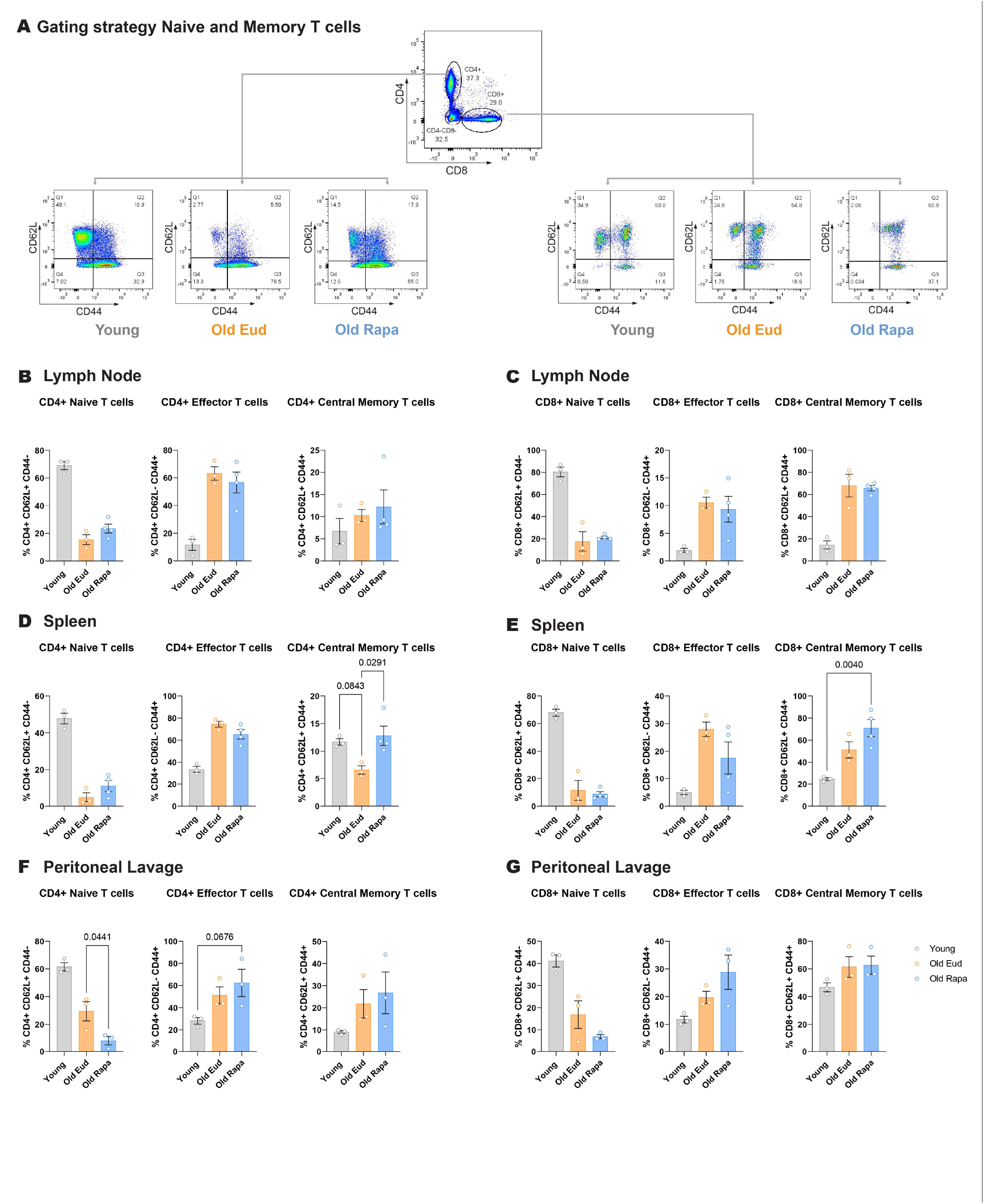
Dietary rapamycin has minor effects on CD4+ and CD8+ T cell populations in aged mice. (**A**) Representative flow cytometry gating strategy for T cells. Gating was performed sequentially on FSC-A versus SSC-A, singlets, live cells, and CD3+ cells, as shown in Fig. 4A (n = 3–4 per group). (**B–G**) Quantification of CD62L and CD44 expression in CD4+ and CD8+ T cells from lymph nodes, spleen, and peritoneal lavage (n = 3–4 per group). Data are presented as mean ± s.e.m. Statistical analysis was performed by One-way ANOVA.

CD4+ T cells isolated from lymph node and spleen of rapamycin-fed mice showed a slight increase in naïve T cells in lymph nodes and spleen, as well as in central memory CD4+ T cells in the spleen, although differences did not reach statistical significance.

No differences were observed for CD8^+^ T cells in the lymph node, albeit an increase was observed in CD8^+^ central memory T cells in the spleen of rapamycin-fed mice compared to both the young and old Eudragit groups (**Fig 3E**). This corroborates previous literature reporting that mTORC1 inhibition via rapamycin can induce an increase in the generation of CD8+ memory T cells (29).

In the peritoneal lavage, CD4+ T cells showed a switch from naïve to effector T cells (**Fig 3F&G**), which appeared specific to the rapamycin treatment. To a certain extent, CD8+ T cells in the peritoneal lavage showed a similar trend, although it did not reach statistical significance.

Overall, these data show modest changes in T cell populations and differentiation to CD8+ T memory cells following rapamycin treatment, in line with previous reports.

### Rapamycin diet induces a decrease in IL-17-producing γδ T cells

While the effect of rapamycin on T cells expressing the main alpha/beta T cell receptor have been investigated, the effect on the less abundant gamma-delta T cell population, which express a type of T cell receptor that does not require MHC recognition, are not well described. In the context of cancer, rapamycin has been shown to enhance γδ T cell responses (30), however, the effect of long-term rapamycin administration on γδ T cell proportion remains unknown. For this reason, we next assessed whether rapamycin diet could have an effect on γδ T cell subsets.

Peritoneal γδ T cells are subdivided in IFN-γ- and IL-17-producing cells, distinguished by the expression of surface marker CD27 (31). CD3+, CD4–, CD8–, γδ TCR+, CD27+ and CD27– cells were assessed in the peritoneal lavage, as well as spleen and lymph node (**Fig 4**).

**Figure 4.**
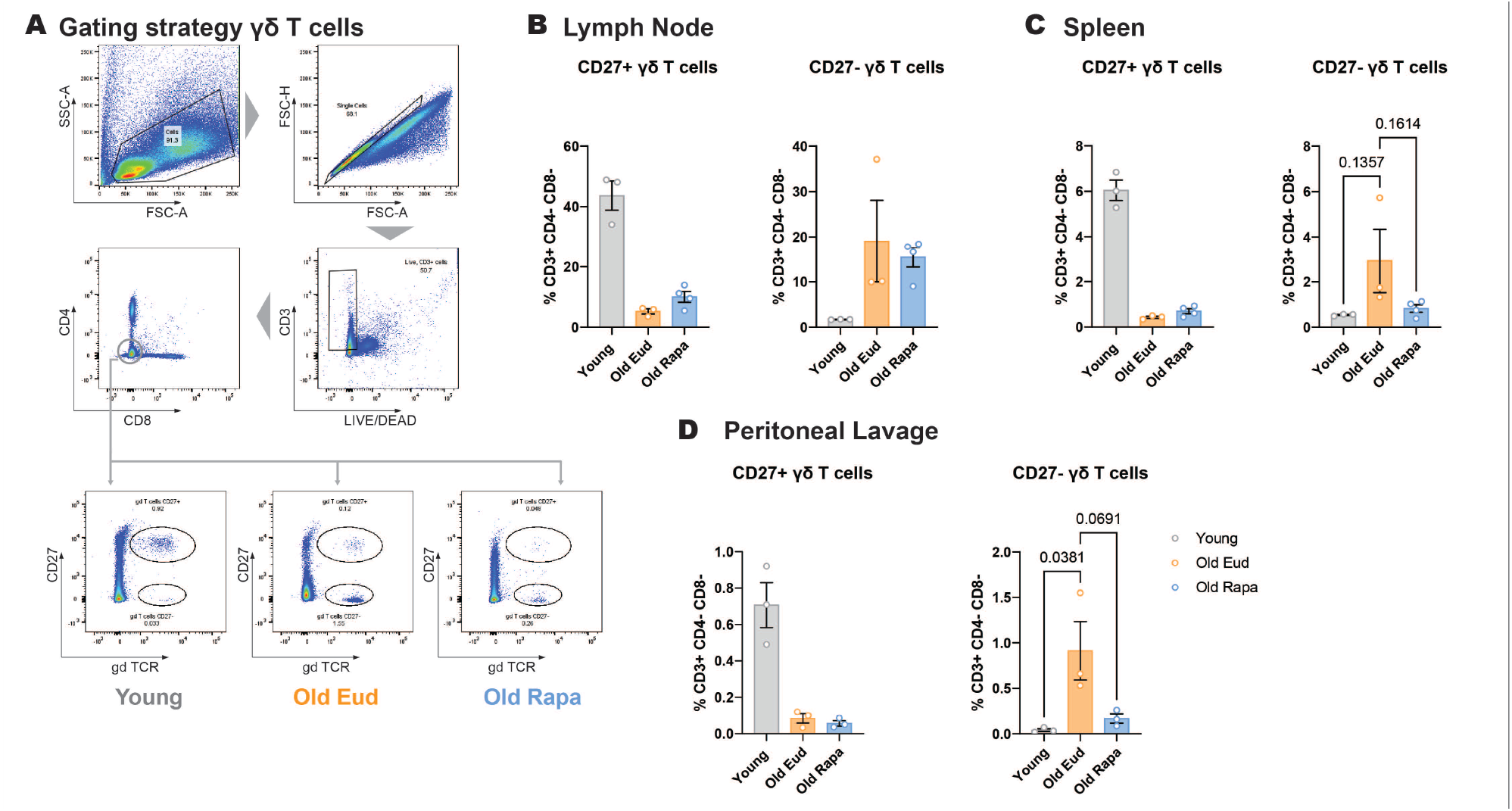
Rapamycin prevents age-associated increase in IL-17-producing γδ T cells. (**A**) Representative flow cytometry gating strategy for identification of γδ T cells. (**B–D**) Quantification of CD27+ and CD27– γδ T cells in the peritoneal lavage, spleen, and lymph nodes of aged mice (n = 3–4 per group). Data are presented as mean ± s.e.m. Statistical analysis was conducted by One-way ANOVA.

In the peritoneal lavage, CD27+ γδ T cells showed a decrease, while CD27– γδ T cells showed an increase with age (**Fig 4A&B**). In contrast, while rapamycin did not affect CD27+ γδ T cells, it significantly affected the IL-17producing, CD27– counterparts, rescuing their levels to those observed in young mice (**Fig 4B**). Similar trends were observed in the spleen and lymph nodes, although differences did not reach statistical significance (**Fig 4C&D**).

Taken together, these data indicate that rapamycin prevents the increase in IL-17-secreting γδ T cells that normally accompanies aging.

### Rapamycin diet modulates microglia reactivity

Our data indicated that long-term rapamycin did not significantly affect the proportion of peripheral immune cells, except for a clear effect on a subset of IL-17-producing γδ T cells.

Given these differences, we investigated whether systemic levels of inflammatory cytokines were also altered (Schematic in **Fig 5A**). To this end, serum cytokine levels were assessed, however, we did not observe any differences in serum concentration of IL-17A together with IL-6, IL-10, IL-27 (**Fig 5B**).

**Figure 5.**
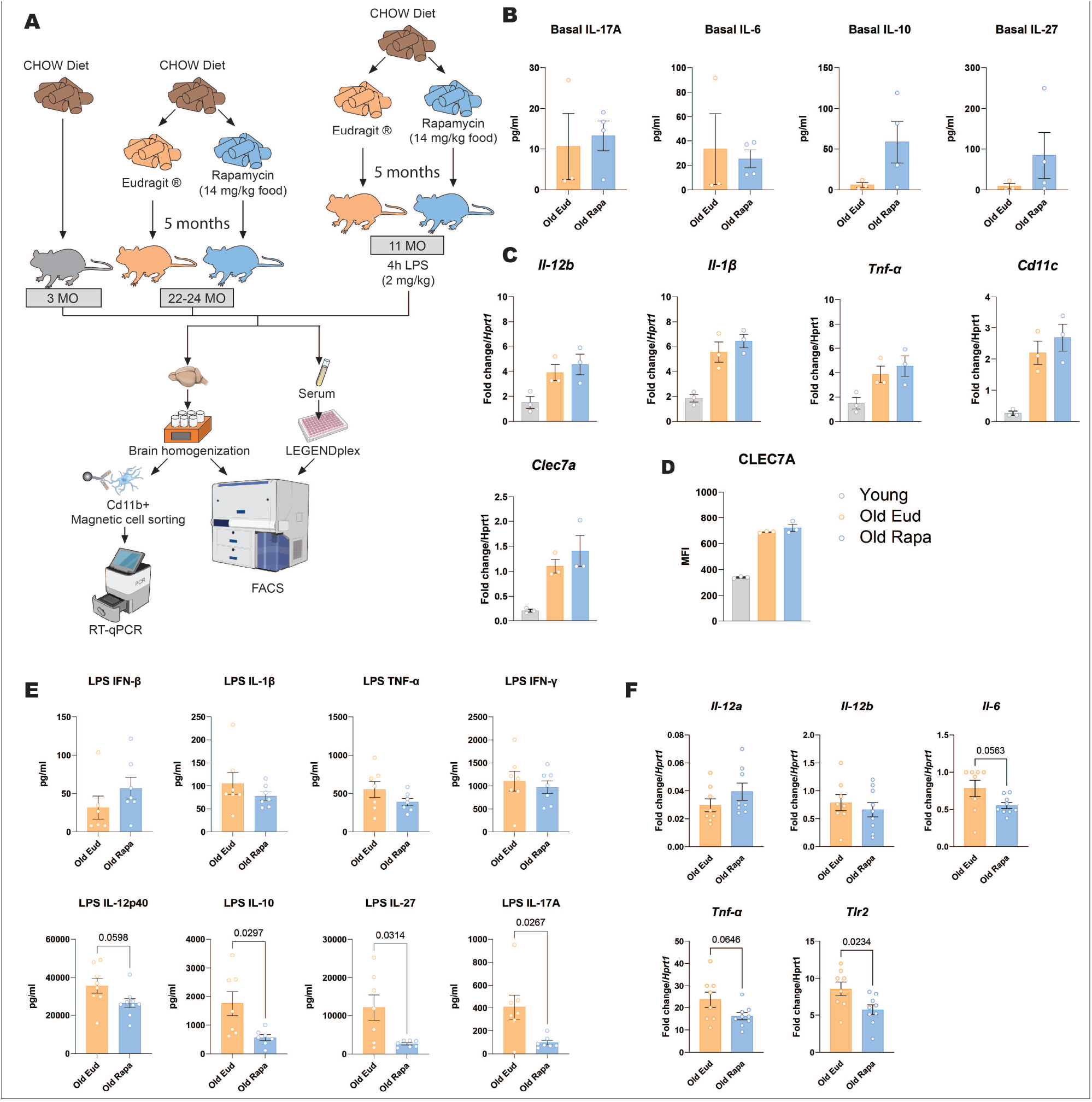
Rapamycin diet reduces inflammation in aged mice challenged with LPS. (**A**) Schematic of young and aged mice fed a control or rapamycin-supplemented diet, with or without LPS stimulation, followed by RT-qPCR and flow cytometry analysis of myeloid immune cells. (**B**) Serum cytokines concentrations in basal conditions. (**C-D**) Basal expression of inflammatory cytokines and receptors in microglia under unstimulated conditions (n = 3 per group). (**E**) Serum cytokine concentrations in LPS-treated mice (n = 7 per group). (**F**) Expression of inflammatory cytokines in microglia following LPS stimulation (n = 8–9 per group). Data are presented as mean ± s.e.m. Statistical analysis was conducted by Welch’s t-test (**B, E, F**) and One-way ANOVA (**C&D**).

Nonetheless, since IL-17 levels can modulate neuroinflammation (32), we next investigated if the rapamycin diet affected the phenotype of central nervous system (CNS) tissue resident immune cells, microglia. To this end, we assessed the expression of pro-inflammatory cytokines and reactive microglial markers using quantitative reverse transcription PCR (RT-qPCR) in microglia cells. Although aging is known to induce upregulation of inflammatory cytokines and receptors in microglia, our data indicate that rapamycin did not counteract age-associated neuroinflammation (**Fig 5C&D**).

Although no changes at steady state were observed, we reasoned that systemic and brain inflammation might be affected in response to a strong inflammatory stimulus, such as intraperitoneal exposure to lipopolysaccharide (LPS).

Thus, a separate cohort of mice were fed with either Eudragit or rapamycin diet from 6 months of age for 5 months, then challenged with 2 mg/kg of LPS intraperitoneally and serum cytokine levels and microglia activation were analyzed 4 hours later.

Serum cytokines IL-1β, TNF-α, IL-23, IL-12p70, IFN-β, and IFN-γ were not different between the two diets (**Fig 5E**). However, the inflammatory cytokines IL-12p40, IL-17A and IL-27 were significantly reduced in rapamycin-fed mice upon LPS stimulation, along with the anti-inflammatory cytokine IL-10 (**Fig 5E**).

Examining the microglial response, we found that rapamycin administration moderately reduced inflammatory markers *Tnf-α, Il6* and *Tlr2* gene expression while others, such as *Il-12a* and *Il-12b* remained unaffected (**Fig 5F**).

Taken together, these results suggest that rapamycin diet can in part reduce systemic and brain inflammation in response to a strong systemic inflammatory stimulus.

## Discussion

Hyperactivation of mTORC1 signaling has been associated with impaired metabolic health and reduced longevity across multiple species, including mice and humans (33-34). The detrimental effects of sustained mTORC1 activity may stem, at least in part, from remodeling of the immune landscape and its contribution to inflammation (35). Accumulating evidence demonstrates that attenuation of mTORC1 activity promotes healthy aging and extends lifespan (36–39). Nonetheless, the long-term consequences of chronic, low-level mTORC1 inhibition on the immune system remain incompletely understood. To address this, we tested the effect of low-dose, long-term rapamycin administration, based on the hypothesis that the resulting mild mTORC1 inhibition could remodel the immune landscape of mice and fine-tune their inflammatory status.

In this study, we investigated how the peripheral immune system and the brain tissue-resident macrophages responded to a rapamycin-containing diet administered in adulthood (6-8months) or later in life (17-19 months). Our findings demonstrate that the treatment was well tolerated and overall safe; and while it did not induce major alterations in immune cell populations, it produced subtle yet significant beneficial effects on immune function in old mice.

We first examined the impact of the rapamycin-containing diet on myeloid cell populations in an aged cohort. Previous studies in adult mice have reported that rapamycin treatment can promote expansion of myeloid cells (40) and reshape their transcriptome under chronic conditions (41). In contrast, in our study of C57BL/6J mice, rapamycin supplementation did not significantly affect the abundance of myeloid cells across multiple compartments. Regarding the lymphoid lineage, likewise we observed minimal effects on the abundance of different subpopulations. However, we observed a downregulation of the CD27–, IL-17-producing γδ T cells, more pronounced in the peritoneal lavage, where they have been implicated in cancer (42) and bowel inflammation (43). The decrease in CD27– γδ T cells could be linked to the role of mTORC1 in Th17 cell differentiation, mediated through the PI3K–mTORC1 axis (44–46).

As rapamycin has been proposed also as an anti-aging agent to preserve brain health and cognition, we were particularly interested in assessing the effect of systemic mTORC1 inhibition on brain-resident microglia. It is known that with age, microglia acquire a primed phenotype, characterized by increased reactivity to an inflammatory stimulus (47). In this study, we demonstrated that rapamycin attenuates the priming phenotype, since in response to a peripheral LPS challenge, the upregulation of inflammatory cytokines and receptors was reduced. Although modest, this reduction could prove beneficial to counteract brain inflammation that accompanies peripheral infections in the elderly (48). Our study did not determine the mechanism underlying the reduction in microglia priming, particularly since rapamycin does not efficiently cross the blood-brain barrier (49–51). However, reduced microglia reactivity is likely either mediated by reduced systemic inflammation or by modulation of the gut-brain axis, driven by rapamycin-induced alterations of the gut microbiome. Notably, even non-oral rapamycin treatment can affect the gut microbial composition (52), a core component of the gut–brain axis that has been identified as a contributor to alterations in microglial behavior (53-55). Indeed, upon LPS, we could observe diminished levels of circulating pro-inflammatory cytokines, such as IL-27 and IL-17. On the other hand, the immunomodulatory cytokine IL-10 was also decreased. This effect might be related to the decrease in inflammatory cytokines inducing a weaker feedback signal that upregulates anti-inflammatory responses (56).

Together, our results provide additional insights into how dietary rapamycin modulates the aged immune system: the absence of major changes in immune cell composition, combined with evidence of beneficial effects on CD27– γδ T cells, microglia and cytokine balance, supports the notion that rapamycin can be administered safely without causing overt immunosuppression. This is particularly relevant given the increasing clinical interest in rapamycin and rapalogs as geroprotective interventions (57, 58). Our data highlight the need to carefully dissect tissue-specific and age-dependent responses to mTORC1 inhibition, and they underscore that the beneficial effects of rapamycin in aging likely arise from subtle recalibrations of immune and inflammatory networks rather than extensive immune remodeling, at least at the dose investigated by us.

Further work is needed to determine whether these modest but significant immune effects translate into improved resilience to age-related pathologies, including neuroinflammation, infections, and systemic inflammation. Recent work suggest that peripheral immune signals may influence innate immune cells in the aged brain: IL-17 produced by γδ T cells may act on IL-17 receptors expressed by microglia and modulate their activation state (59). It has recently been shown that gut-derived γδ T17 cells migrate to the meninges, where IL-17A can alter the mitochondrial function of microglia and trigger downstream inflammatory pathways leading to microglial synaptic pruning in mouse models of encephalopathy (60). This raises the intriguing possibility that dietary rapamycin may exert some of its effects on brain microglia indirectly via modulation of peripheral γδ T cell responses, although further work is needed to test this hypothesis in the context of aging. In summary, our findings contribute to a growing body of evidence suggesting that dietary rapamycin is a safe and promising strategy for promoting healthy aging by modulating the aged immune system.

## Methods

### Mice

C57BL6J female and male mice of different ages, as indicated in figure legends, were purchased from Charles River and fed a chow rapamycin-containing diet for a period of 5 months. Mice were housed at Barts Cancer Institute, Queen Mary University of London, UK and DZNE Bonn, Germany and experiments were carried out under UK Home Office PPL 70/7411 and German TVA 81-02.04.2018.A215.

### Rapamycin diet

Food-encapsulated rapamycin was prepared following the protocol by Harrison *et al*., (18).

Rapamycin was obtained from Rapamycin Holdings (now Entomora Biosciences) in San Antonio, Texas, USA. Rapamycin was microencapsulated with an enteric coating material, Eudragit S100 (Röhm Pharma, Germany), which protects rapamycin through the food preparation process. Since Eudragit S100 is water soluble only at non-acidic pH, rapamycin is only released in the small intestine and protected in the stomach. Eudragit-microencapsulated rapamycin and Eudragit as control were incorporated into separate LabDiets on a 5058 base by Purina Labs. The amount of rapamycin used corresponded to 152 ppm, resulting in a 14 ppm concentration of active rapamycin in mice (2.24 mg of rapamycin per kg body weight).

### In vivo LPS injection

Female 6-month-old mice were fed a rapamycin- or Eudragit-diet for 5 months, then injected with 2 mg/kg of LPS of E coli 0111:B4 (Sigma, Cat. No. 297-473-0) in 200 μL PBS or PBS only as a control. Intraperitoneal injections were performed using a 27G needle and 1 mL syringe. Mice were sacrificed after 4 hours and organs were collected as described below.

### Organ processing

#### Blood

Blood was obtained by cardiac puncture using a 27G needle and 1 mL syringe coated in 0.5 M Ethylenediaminetetraacetic acid (EDTA). It was then transferred into a 1.5 mL EDTA-coated tube and placed on ice before being centrifuged 3 times at increasing speed, removing pellets each time. Centrifugation speeds were 200 x g for 3 min; 2,300 x g for 5 min and 16,100 x g for 3 min. Plasma was then aliquoted and snap frozen before being stored at -80°C until cytokine assessment was carried out.

#### Spleen

Spleens were harvested, placed in 10 mL of PBS and kept on ice until further processing. Afterwards, they were placed on a 70 μm cell strainer from Fisher Scientific (Cat. No. 11597522). They were gently mashed through the strainer using a 5 mL syringe plunger. 10 mL of PBS was added to the cell suspension before centrifugation at 300 x g for 10 minutes at 4°C.

#### Bone marrow

Cells were obtained from the bone marrow by flushing the femurs and tibia with Dulbecco’s Modified Eagle Medium (DMEM, Sigma, D5768) supplemented with 10% Fetal Bovine Serum (FBS, Life Technology, Cat. No. 10500-064) and 1% penicillin/streptomycin (Sigma, Cat. No. P4333). 1 mL of complete media was flushed through the bone marrow using a 27G needle, several times. Cells were then strained through a 70 μm cell strainer before 20 mL of PBS was added to the cell suspension and cells were spun at 300 x g for 10 minutes 4°C.

#### Brain

The brain was dissected following perfusion with 35 mL of ice-cold Hank’s balanced salt solution (HBSS) for 5 minutes and subsequently digested using a neuronal dissociation kit (Miltenyi Biotec, Cat. No. 130-092-628). Next, cells were washed twice in HBSS and myelin was removed by negative selection using myelin removal beads (Miltenyi Biotec, 130-094-060) in MACS LS columns (Miltenyi Biotec, Cat. No. 140-096-433). Briefly, 1.8 mL of fluorescence-activated cell sorting (FACS) buffer [phosphate buffered saline (PBS) + 0.5% bovine serum albumin (BSA) + 2mM EDTA] and 200 µL of myelin removal beads were added to each brain cell suspension and incubated at 4°C in the dark for 15 minutes. Following this incubation, 18 mL of FACS buffer was added to wash the cells, then centrifuged for 10 minutes at 300 x g. Using three LS columns per brain, 1 of 3 mL of beads/cell suspension were loaded onto each column placed in a QuadroMACS separator magnet (Miltenyi Biotec, Cat No. 130-042-302). This step was required to separate bead-bound myelin from the cell suspension. Each LS column was washed twice with FACS buffer and each time the flow-through was collected in order to collect brain cells.

### Cell processing for flow cytometry

Cells from blood, spleen and bone marrow were first subjected to red blood cell lysis using 10X red blood cell lysis buffer (eBioscience, Cat. No. 00-4300-54), diluted 1:10 in distilled water. Cell pellets were resuspended in 10 mL of 1X lysis buffer and placed in the dark for 10 minutes at room temperature. Following this incubation, cells were washed twice with FACSbuffer [phosphate buffered saline (PBS) + 0.5% bovine serum albumin (BSA) + 2mM EDTA] and used for subsequent antibody staining.

### Flow cytometry staining

Cells were stained in 96-well V-bottom plates from Thermo Fisher (Cat.No. 2605), 2 x 10^6 cells in 100 μL were added to each well and the plate was centrifuged at 300 x g for 5 minutes at 4°C. Cells were then resuspended in 30 μL of CD16/CD32 FcR block (eBioscience, Cat. No. 14-0161-86) for 15 minutes at 4°C. Following FcR block, a 2X antibody mix was prepared using FACS buffer and 30 μL was added to each well. All relevant information regarding the panel of antibodies used (manufacturer, clone and dilutions) is summarized in Tab. 1. After a 30 min incubation at 4°C in the dark, cells were washed twice with 150 μL of FACS buffer before being resuspended in 150 μL of FACS buffer, which was transferred to 1.2 mL FACS tubes (Star lab, Cat. No. I1412-7400). Data acquisition was carried out with a 4-laser LSRFortessa or a 5-laser FACSymphony (BD Biosciences) and data analysis was done using FlowJo software (BD Bioscience).

**Tab. 1.** List of antibodies and fluorescent dyes implemented in flow cytometry experiments.

| Target | Manufacturer | Cat Number | Clone | Dilution factor |
| --- | --- | --- | --- | --- |
| CD4 | eBioscience | 17-0041-83 | GK1.5 | 1:100 |
| CD8 | eBioscience | 46-0081-80 | 53-6.7 | 1:100 |
| CD45 | eBioscience | 9047-9459-120 | 2D1 | 1:100 |
| CD62L | eBioscience | 12-0621-85 | MEL-14 | 1:100 |
| CD11B | eBioscience | 25-0112-82 | M1/70 | 1:200 |
| LY6C | eBioscience | 48-5932-82 | HK1.4 | 1:100 |
| LY6G | eBioscience | 17-5931-81 | RB6-8C5 | 1:100 |
| F4/80 | eBioscience | 11-4801-85 | BM8 | 1:100 |
| CLEC7A | eBioscience | 17-5859-80 | Bg1fpj | 1:100 |
| $\gamma\delta$ TCR | Biolegend | 118108 | GL3 | 1:100 |
| CD27 | Biolegend | 124216 | LG.3A10 | 1:100 |
| TLR2 | eBioscience | 12-9021-80 | 6C2 | 1:100 |
| PS6 | Cell Signaling | 5018 | N/A | 1:50 |
| DAPI | Cambridge Bioscience | 40043-BT | N/A | 2.5 $\mu\text{g/mL}$ |
| FIXABLE VIABILITY DYE (FVD) | eBioscience | 65-0863-18 | N/A | 1:200 |

### Microglia isolation

Following brain homogenization (see Organ harvesting: Brain) single cell suspension was resuspended in 500 μL of FACS buffer and 40 μL of CD11b microbeads (Miltenyi Biotec, Cat. No. 130-093-636) and incubated at 4°C in the dark for 15 minutes. Following this incubation, cells were washed with 3 mL of FACS buffer and centrifuged for 10 min at 300 x g at 4°C. Using one MS column (Miltenyi Biotec, Cat. No. 130-042-201) per sample, 500 μL of bead/cells were loaded onto each column placed in an OctoMACS separator magnet (Miltenyi Biotec, Cat. No. 130-042-108). Each MS column was washed twice with FACS buffer and then 1 mL of FACS buffer was used to flush CD11b-positive cells using the plunger provided. Purity of microglia was assessed by flow cytometry analysis by staining an aliquot of cells with the microglia-specific markers CD11b and CD45 and reached 92-95%.

### RNA isolation

Microglia (0.5-1 x 10^6) were lysed in 350 μL RLT lysis buffer (Qiagen, Cat. No. 74004) and stored at -80°C. RNA was extracted with the RNeasy microkit from Qiagen (Cat. No. 74004), according to the manufacturer guidelines.

### cDNA synthesis and Quantitative Real-Time Polymerase Chain Reaction (qPCR)

Following RNA extraction, RNA was reverse transcribed into cDNA using the High-Capacity cDNA Reverse Transcription kit from Applied Biosystems (Cat. No. 436-8814). Briefly, a master mix was prepared that contained (per reaction): 1X RT buffer, 1X random hexamers, 4 mM dNTP, 1 μL of reverse transcriptase (50 U/ml) and 2.2 μL of nuclease-free water. This master mix was added to the extracted RNA, contained in 12 μL, for a final volume per reaction of 20 μL. All samples were then incubated in a thermocycler using the following program; 25°C for 10 minutes, 37°C for 120 minutes, 85°C for 5 minutes and held at 4°C. Following cDNA synthesis, 130 μL of nuclease-free water was added to each sample to dilute the enzymes used in the cDNA synthesis. A master mix was then prepared for the qPCR for each gene of interest, which included per reaction: 10 μL of Biorad iTaq master mix (Biorad, Cat. No. 436-814) and 1 μL of each FAM labelled gene of interest. This master mix was added to a qPCR plate and 9 μL of diluted cDNA was added to each corresponding well. The Step One Plus Real Time PCR machine from Applied Biosystems was used to carry out the qPCR. The PCR program included an initial denaturation cycle that lasted for 10 minutes at 95°C, followed by 40 cycles of amplification that included 15 seconds at 95°C and one minute at 60°C; the last part of the program included one more cycle at 25°C for 15 seconds. All primers used are summarized in Tab. 2.

**Tab. 2.** List of primers implemented in Real Time qPCR analysis.

| Target gene | Manufacturer | Cat Number |
| --- | --- | --- |
| *Clec7a* | Applied Biosystems | Mm01183349_m1 |
| *Tlr2* | Applied Biosystems | Mm00442346_m1 |
| *Tnf-α* | Applied Biosystems | Mm00443258_m1 |
| *Il-1b* | Applied Biosystems | Mm00434228_m1 |
| *Il-12b* | Applied Biosystems | Mm01288989_m1 |

### LEGENDplex

Cytokine levels in serum or in microglia protein lysates were determined using the Legendplex mouse inflammation 13-plex panel, (Biolegend, Cat. No. 740446); a bead-based assay that allows simultaneous measurement of analytes based on cell size using flow cytometry. Specifically, the cytokines assessed were IFN-β, IFN-γ, IL-1β, IL-6, IL-10, IL-12B, IL-17A, IL-27 and TNF-α. The assay was carried out in a 96-well plate following manufacturers’ instructions. For TNF and IL-6 measurements on microglia culture supernatants, the BD OptEIA ELISA kits were used (BD Pharmingen, Cat. No. 558534, 555240) and the assay was carried out according to manufacturers’ guidelines.

### ELISA

ELISA kits were obtained from BD Biosciences: IL-12p40 (Cat. No. 555165), IL-6 (Cat. No. 555240) and TNFα (Cat. No. 558534). ELISA was performed according to the manufacturer’s guidelines; however, half-surface area 96-well plates were obtained from Corning Costar (Cat. No. 3690) and used for the assay. The wells in these plates are half the volume of standard 96-well plates, allowing the use of a reduced number of samples and reagents. This was particularly important for plasma cytokine levels, as the amount of plasma obtained from each mouse was potentially a limiting factor. Briefly, capture antibody was diluted in PBS and added to each well of a 96-well plate. The plate was then incubated overnight at 4°C, prior to being washed three times with wash buffer (PBS with 0.05% Tween-20). The plate was then incubated with assay diluent (PBS with 10% FBS) for 1 hour, in order to block non-specific binding and washed three times with wash buffer. The higher concentration of standards (1000 pg/mL) were made using a stock solution of each cytokine and then a serial dilution was performed on a separate plate to obtain the remaining standards (500, 250, 125, 62.5, 31.3 and 15.6 pg/mL). Samples and standards were then added to the ELISA plate and incubated for 2 hours at room temperature. Samples that required dilution were diluted in assay diluent. Following incubation, the plate was washed 5 times with wash buffer and then the working detector solution (detection antibody and HRP) was added to the plate and it was incubated at room temperature for 1 hour. The plate was then washed 7 times before the substrate solution [Tetramethylbenzidine (TMB) and hydrogen peroxide from the TMB Substrate Reagent Set (BD PHarmingen, Cat. No.555214)] was added and incubated in the dark for a maximum of 30 minutes. Following the appropriate color change, 25 ml of 1M H3PO4 was added to stop the reaction. Plates were analyzed on a plate reader plate reader to read the optical density at 450 nm.

### Statistical analysis

Statistical analysis was conducted used GraphPad Prism, one-way ANOVA and Welch’s t-test were employed as required and as reported in the respective figure legends. A p value < 0.05 was considered statistically significant. P values below 0.1 are reported in the figures.

## Acknowledgements

We thank Profs Frances Balkwill and Federica Marelli-Berg (Queen Mary University of London) and Dr Dan Ehninger (DZNE, Bonn) for useful discussions. We also thank previous and current members of the Capasso group for experimental support.

